# Active sensing with light improves predator detection in a diurnal fish

**DOI:** 10.1101/324202

**Authors:** Matteo Santon, Pierre-Paul Bitton, Jasha Dehm, Roland Fritsch, Ulrike K. Harant, Nils Anthes, Nico K. Michiels

## Abstract

Active sensing by means of light is rare. In vertebrates, it is known only from chemiluminescent fish with light organs below their pupils, an anatomical arrangement that is ideal to generate eyeshine in the pupils of nearby organisms. Here, we test whether diurnal fish can achieve the same by redirecting sunlight through reflection instead. We recently showed that small (< 5 cm), benthic, marine triplefin fish actively redirect downwelling light using their iris. We hypothesized that this mechanism allows triplefins to improve detection of a cryptic organism by generating eyeshine in its pupil. Here, we tested this by attaching small dark hats to triplefins to shade their iris from downwelling light. Two controls consisted of triplefins with a clear or no hat. These treatments test the prediction that light redirection increases the visual detection ability of triplefins. To this end, we placed treated fish in a tank with a display compartment containing either a stone as the control stimulus, or a scorpionfish, i.e. a cryptic, motionless triplefin predator with retroreflective eyes. After overnight acclimatization, we determined the average distance triplefins kept from the display compartment over two days. Both in the laboratory (*n* = 15 replicates per treatment) and in a similar field experiment at 15 m depth (*n* = 43 replicates per treatment) fish kept longer distances from the scorpionfish than from the stone. This response varied between hat treatments: shaded triplefins stayed significantly closer to the scorpionfish in the laboratory and in one of two orientations tested in the field. A follow-up field experiment at 10 m depth revealed the immediate response of triplefins to a scorpionfish. At first, many individuals (*n* = 80) moved towards it, with shaded triplefins getting significantly closer. All individuals then gradually moved to a safer distance at the opposite half of the tank. Visual modelling supported the experimental results by showing that triplefins can redirect enough light with their iris to increase a scorpionfish’s pupil brightness above detection threshold at a distance of 7 cm under average field conditions and at more than 12 cm under favorable conditions. We conclude that triplefins are generally good in the visual detection of a cryptic predator, but can significantly improve this ability when able to redirect downwelling light with their iris and induce eyeshine in the predator’s pupil. We discuss the consequences of “diurnal active photolocation” for visual detection and camouflage among fish species.

## Introduction

The only vertebrates known to use light for active sensing are nocturnal and deep-sea fish with a subocular chemiluminescent light organ [1–3]. Recent findings in the triplefin *Tripterygion delaisi* suggested that diurnal fish may use an analogous mechanism that exploits downwelling sunlight and redirects it sideways using the iris, generating a phenomenon called “ocular spark” (Fig. 1a-b)[4]. Ocular sparks can arise because in fish the lens usually protrudes from the pupil. This allows downwelling light to cross the lens and be focused on the iris below. This process can be controlled by subtle eye movement [4]. The resultant bright focal point reflects sunlight sideways outside the range dictated by Snell’s window, which constrains downwelling sunrays to a 96° cone pointing down from the surface [5]. The authors hypothesized that ocular sparks may be sufficient to illuminate the immediate surroundings and improve visual detection of cryptic organisms, a process called “diurnal active photolocation”. Because the absolute amount of redirected light is small, the structures that can be detected in this way can be predicted to be nearby and highly reflective. We therefore focus on retroreflective eyes, which are among the strongest directional reflectors found in nature. Their key properties are a focusing lens in front of a reflective layer [6, 7]. This design is known to improve dim light vision [8], but in some cryptic species also enhances camouflage of an otherwise conspicuously black pupil during the day [9, 10] (Fig. 1d-f). As a side-effect, however, retroreflective eyes can be easily revealed when illuminated with a source next to the observer’s eye. This specific configuration is required, as the retroreflected light is returned towards the source in a narrow beam [11, 12]. When this coaxial alignment of light source and detecting eye is given, even weak illumination can generate eyeshine in a nearby retroreflective target (Fig. 1e, video clip in supplement of [4]). This is also the accepted explanation for why the light organ of chemiluminescent fishes is located just below their pupil [2]. Yet, it remains to be demonstrated whether light redirection by triplefins can work in a similar way [4, 13].

**Figure 1.**
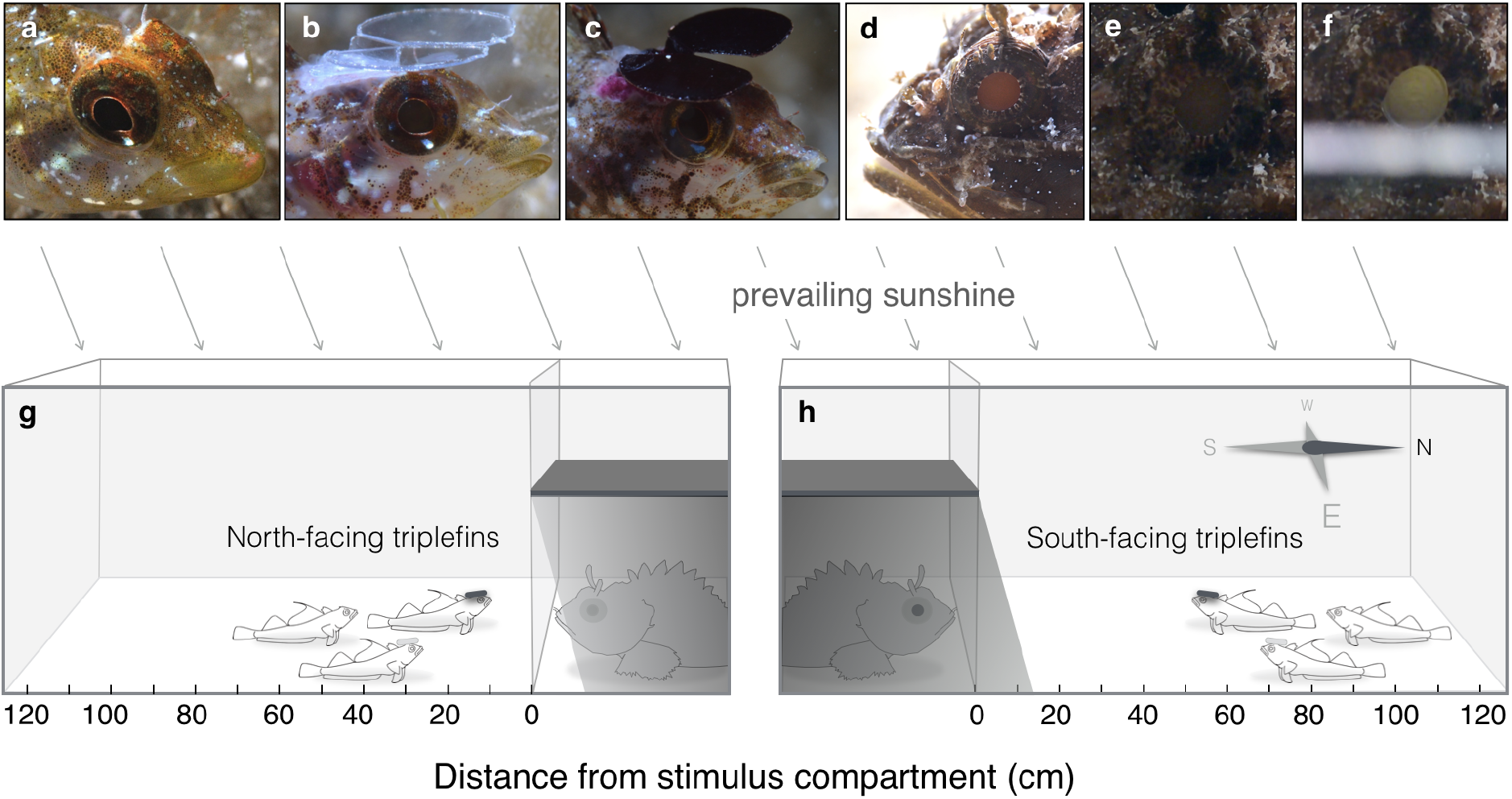
Overview of experimental manipulation and design. Triplefins (*Tripterygion delaisi*) were subjected to one of three treatments: **a.** Unhatted sham control, **b.** Clear-hatted control, and **c.** Shading hat treatment. While **a** and **b** can re-direct light using blue ocular sparks (bright bluish dots on the lower iris), **c** cannot. **d.** Scorpionfish (*Scorpaena porcus*) show retroreflective eyeshine [9] when illuminated coaxially, here by means of a strip of matt white paper (**e** and **f**). **g-h.** Triplets of triplefins, one of each treatment, were exposed to a shaded predator or stone (not shown) behind a windowpane. We tested two opposite orientations in the field (triplefins facing north or south). This was not required in the laboratory (not shown). The response variable was distance from the stimulus, measured the day after adding fish to the tanks. Drawings not to scale, see Materials and Methods for dimensions. Pictures by M.S. and N.K.M.

Here, we tested whether ocular sparks improve the ability of triplefins to detect scorpionfish [9], which are common, cryptic, sit-and-wait predators with large pupils and daytime retroreflective eyeshine [9, 14]. To suppress ocular spark generation in triplefins, we glued opaque mini-hats on their heads (Fig. 1c). Two controls permitted unobstructed ocular spark formation: a clear-hatted (Fig. 1b) and an unhatted sham control (Fig. 1a). Triplefins were placed in large tanks and shown one of two visual stimuli placed in the shade behind a windowpane: a scorpionfish or a stone. We expected triplefins to be attracted to the display compartment as they prefer hard substrates with shady edges over the shade-free sand in their own compartment. However, we also expected them to keep a safe distance after recognizing the scorpionfish. We predicted that shaded triplefins, deprived of the ability to use active photolocation, would display shorter “safe distances” from a scorpionfish compared to the controls. No such effect was expected for the stone stimulus. We tested this paradigm independently in the laboratory and in a field setup at 15 m depth. In both experiments, we used triplets consisting of one individual from each of the three hat treatments and observed them over two days. In a follow-up field experiment at 10 m, we tested hatted triplefins individually and observed how close they approached a scorpionfish immediately after release. We then monitored their position relative to the scorpionfish during the next 90-100 min. Although these experiments tested the effect of the triplefin’s ability to redirect ambient light, they did not directly test whether the observed effects were caused by an ability to generate eyeshine in the scorpionfish. Using visual modelling, we therefore estimated the distances at which a triplefin can perceive an increase in the brightness of a scorpionfish pupil induced by an ocular spark.

Triplefins are particularly suitable for this type of research. Unlike other small benthic fish such as blennies and gobies, they do not have a hiding place or nest where they spend most of their time [15]. Instead, they roam on the substrate looking for micro-prey. This is made possible by their cryptic coloration [16], their habit of moving cautiously and secretively while assessing their surroundings with independent eye movement and by their high visual acuity and contrast sensitivity [17, 18]. This makes them a convenient system for laboratory and field experiments that include unusual treatments such as hats.

## Results

### Distance from scorpionfish or stone in the laboratory

We recorded the position of each individual triplefin relative to the visual stimulus five times per day over 2 days after triplefins had been acclimatized to their tank for more than 12 h. Due to premature hat loss, 15 out of 20 triplets were available for analysis. Triplefins kept a significantly greater mean distance from the predator than from the stone irrespective of the hat treatment (Figure 2), indicating that vision alone already allowed detection of the scorpionfish independent of diurnal active photolocation. This effect was indistinguishable between the clear-hatted and unhatted controls (Table 1a), showing that the hat manipulation did not affect fish behavior. For subsequent comparisons, the controls were thus averaged per triplet and observation. A comparison of the distances measured in controls relative to the shading hat treatment (Figure 2, Table 1b) confirmed the overall effect of the stimulus, but included an effect of hat treatment. Relative to the controls, shaded individuals stayed significantly closer to the scorpionfish (Table 1c). This was not the case when exposed to the stone (LMEM for stimulus stone: hat treatment *p* = 0.21). The predictor variable *time of day* did not contribute significantly to the model, indicating that movements towards or away from the stimulus were balanced during the observation period.

**Table 1.** Statistical analysis of the laboratory data presented in Figure 2. Generalized Linear Mixed Models with distance from the two visual stimuli (scorpionfish or stone) as the response variable. Given that the two control treatments did not differ in their response to the two stimuli (**a**), their respective measurements were averaged for the main analysis (see Fig. 2) that compared the response of control and shaded treatments to both stimuli (**b**, Figure 2). The final model (**c**) tests the difference between the controls and the shaded treatment in their response to the scorpionfish only. CI = credible interval. For factorial predictors, estimates are computed using the indicated intercept levels as reference. This choice is arbitrary and does not affect overall conclusions.

| Predictors | Predicted mean | Lower 95% CI | Upper 95% CI | P |
| --- | --- | --- | --- | --- |
| **a. Response of unhatted and clear-hatted controls to both stimuli**<br>*n* = 15 triplets, $R^2_{\text{marg}} = 0.30$, $R^2_{\text{cond}} = 0.31$ | | | | |
| *Intercept* (stone & no hat) | 25.770 | 16.031 | 35.524 | < 0.0001 |
| *Treatment* (clear hat) | 3.416 | -4.110 | 10.881 | 0.373 |
| *Stimulus* (scorpionfish) | 32.917 | 25.385 | 40.419 | < 0.0001 |
| *Treatment x Stimulus* | -4.421 | -14.913 | 6.227 | 0.412 |
| *Stimulus order* | 4.526 | -0.791 | 9.886 | 0.100 |
| **b. Response of averaged controls and shaded individuals to both stimuli**<br>*n* = 15 triplets, $R^2_{\text{marg}} = 0.28$, $R^2_{\text{cond}} = 0.28$ | | | | |
| *Intercept* (stone & controls) | 31.994 | 27.466 | 36.667 | < 0.0001 |
| *Treatment* (shading hat) | -4.849 | -11.388 | 1.683 | 0.151 |
| *Stimulus* (scorpionfish) | 30.700 | 25.304 | 35.980 | < 0.0001 |
| *Treatment x Stimulus* | -11.390 | -20.570 | -2.142 | 0.017 |
| *Stimulus order* | 4.580 | 0.190 | 8.936 | 0.041 |
| **c. Response of averaged controls and shaded individuals to the scorpionfish stimulus only**<br>*n* = 15 triplets, $R^2_{\text{marg}} = 0.14$, $R^2_{\text{cond}} = 0.23$ | | | | |
| *Intercept* (controls) | 62.918 | 57.127 | 68.660 | < 0.0001 |
| *Treatment* (shading hat) | -16.220 | -21.417 | -11.043 | < 0.0001 |
| *Stimulus order* | 4.256 | -4.007 | 12.420 | 0.331 |

**Figure 2.**
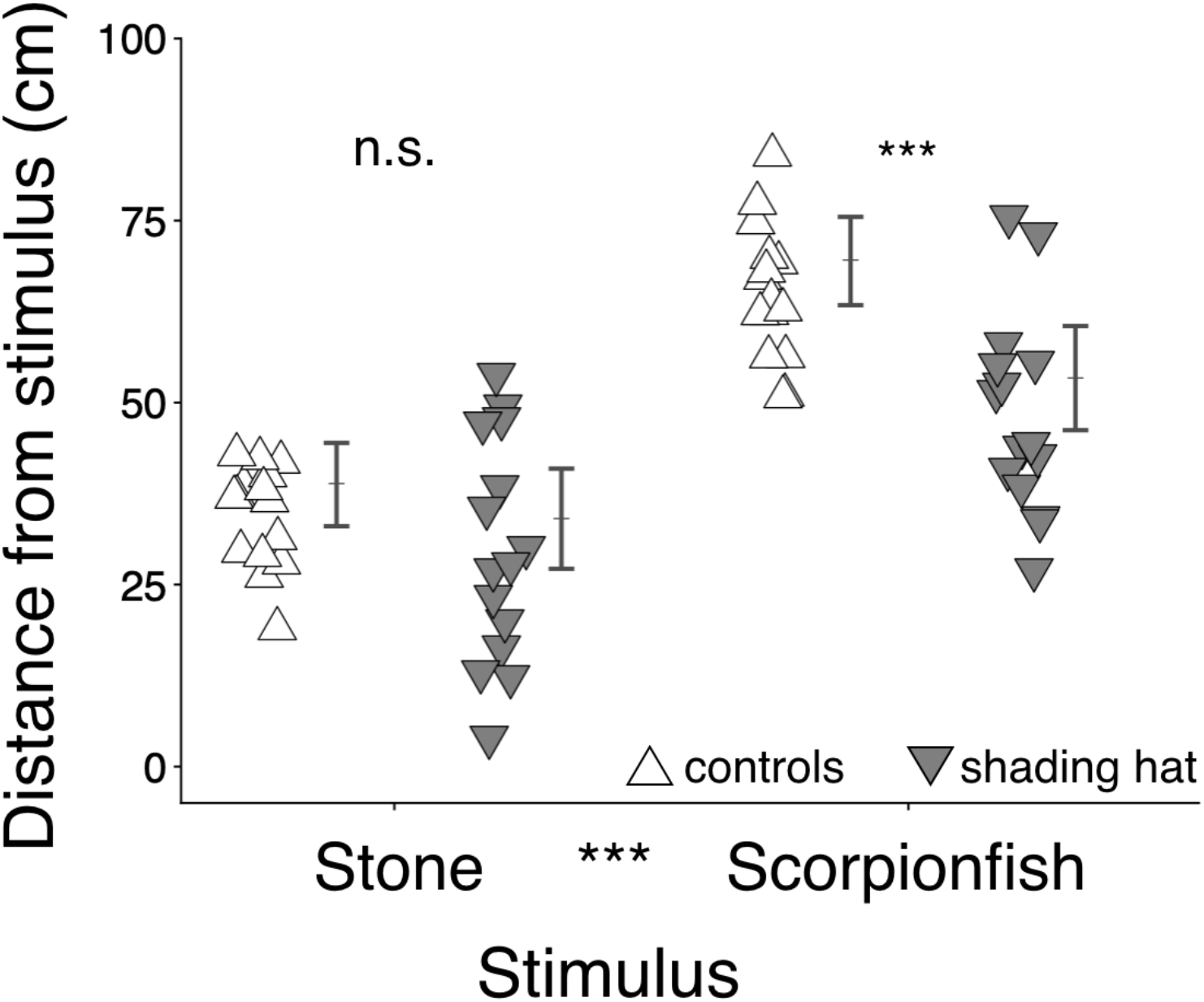
Consequences of hatting in the laboratory shown as the average distance from the stimulus compartment as a function of stimulus type (stone or scorpionfish) and hat treatment. Relative to the controls, shaded individuals stayed significantly closer to the scorpionfish. Symbols = average of 5 measurements per triplet; *n* = 15 triplets; error bars: model-predicted group means ± 95 % credible intervals; *** = *p* < 0.001, n.s. = *p* > 0.05 (see Table 1 and Methods). Note that statistical comparisons between treatments rested on the connected measures *within* triplets and 5 observations per stimulus, making group means and error bars imprecise indicators of the statistical significance of paired measures.

### Distance from scorpionfish or stone in the field

We replicated the experiment in 10 transparent tanks on the sea floor at 15 m depth (Figure 1g-h). Anticipating an effect of orientation relative to the sun without *a priori* expectation, five tanks were oriented north, another five south (Figure 1h). We recorded the distance of each individual to the stimulus compartment during three dives in the course of a day, after triplefins had been acclimatized to their tank for more than 12 h. Forty-three triplets were available for the final statistical analysis. In agreement with the laboratory experiment (Table 1a), the two control treatments kept very similar distances from each combination of stimulus and orientation (Table 2a). However, south-facing controls responded stronger to the scorpionfish than north-facing controls, resulting in a significant stimulus x orientation interaction (Table 2a). For subsequent comparisons, the controls were again averaged per triplet, and the analyses performed separately for the two tank orientations.

**Table 2.**
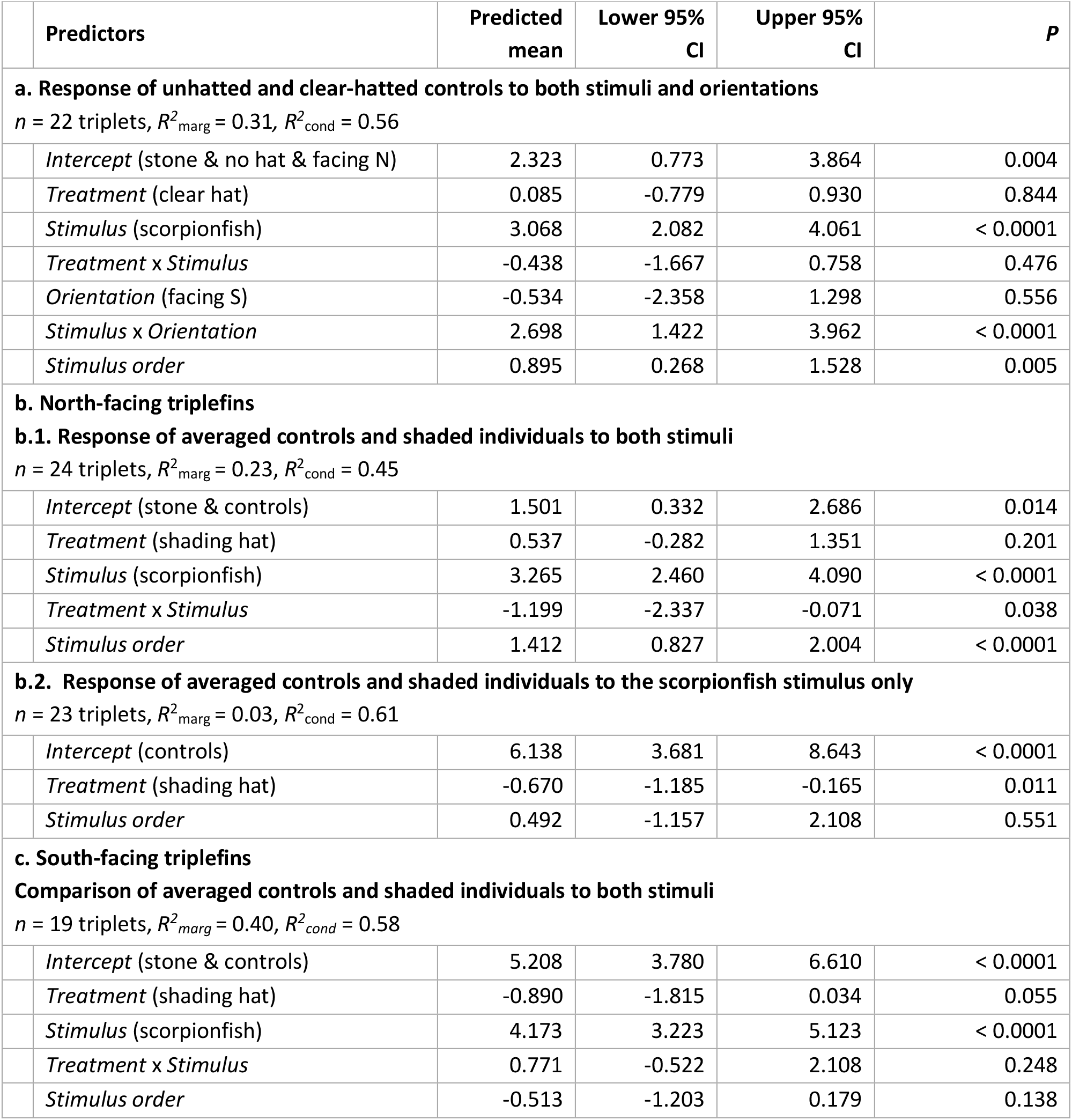
Statistical analysis of the field data presented in Figure 3. Generalized Linear Mixed Models with the distance from the two visual stimuli (scorpionfish or stone) as the response variable. Given that the two control treatments did not differ in their response to the two stimuli (treatment x stimulus interaction in **a**), the respective measurements were averaged for the main analysis (see Fig. 3) that compared the response of control and shaded treatments to both stimuli split by the two orientations (**b-c**, Figure 3). Predicted means and their credible intervals (CI) are based on a square-root transformation of the response variable (see Materials and Methods). For factorial predictors, estimates are computed using the indicated intercept levels as reference. This choice is arbitrary and does not affect overall conclusions.

| Predictors | Predicted mean | Lower 95% CI | Upper 95% CI | P |
| --- | --- | --- | --- | --- |
| **a. Response of unhatted and clear-hatted controls to both stimuli and orientations** |  |  |  |  |
| *n* = 22 triplets, $R^2_{\text{marg}} = 0.31$, $R^2_{\text{cond}} = 0.56$ | | | | |
| *Intercept* (stone & no hat & facing N) | 2.323 | 0.773 | 3.864 | 0.004 |
| *Treatment* (clear hat) | 0.085 | -0.779 | 0.930 | 0.844 |
| *Stimulus* (scorpionfish) | 3.068 | 2.082 | 4.061 | < 0.0001 |
| *Treatment* x *Stimulus* | -0.438 | -1.667 | 0.758 | 0.476 |
| *Orientation* (facing S) | -0.534 | -2.358 | 1.298 | 0.556 |
| *Stimulus* x *Orientation* | 2.698 | 1.422 | 3.962 | < 0.0001 |
| *Stimulus order* | 0.895 | 0.268 | 1.528 | 0.005 |
| **b. North-facing triplefins** |  |  |  |  |
| **b.1. Response of averaged controls and shaded individuals to both stimuli** |  |  |  |  |
| *n* = 24 triplets, $R^2_{\text{marg}} = 0.23$, $R^2_{\text{cond}} = 0.45$ | | | | |
| *Intercept* (stone & controls) | 1.501 | 0.332 | 2.686 | 0.014 |
| *Treatment* (shading hat) | 0.537 | -0.282 | 1.351 | 0.201 |
| *Stimulus* (scorpionfish) | 3.265 | 2.460 | 4.090 | < 0.0001 |
| *Treatment* x *Stimulus* | -1.199 | -2.337 | -0.071 | 0.038 |
| *Stimulus order* | 1.412 | 0.827 | 2.004 | < 0.0001 |
| **b.2. Response of averaged controls and shaded individuals to the scorpionfish stimulus only** |  |  |  |  |
| *n* = 23 triplets, $R^2_{\text{marg}} = 0.03$, $R^2_{\text{cond}} = 0.61$ | | | | |
| *Intercept* (controls) | 6.138 | 3.681 | 8.643 | < 0.0001 |
| *Treatment* (shading hat) | -0.670 | -1.185 | -0.165 | 0.011 |
| *Stimulus order* | 0.492 | -1.157 | 2.108 | 0.551 |
| **c. South-facing triplefins** |  |  |  |  |
| **Comparison of averaged controls and shaded individuals to both stimuli** |  |  |  |  |
| *n* = 19 triplets, $R^2_{\text{marg}} = 0.40$, $R^2_{\text{cond}} = 0.58$ | | | | |
| *Intercept* (stone & controls) | 5.208 | 3.780 | 6.610 | < 0.0001 |
| *Treatment* (shading hat) | -0.890 | -1.815 | 0.034 | 0.055 |
| *Stimulus* (scorpionfish) | 4.173 | 3.223 | 5.123 | < 0.0001 |
| *Treatment* x *Stimulus* | 0.771 | -0.522 | 2.108 | 0.248 |
| *Stimulus order* | -0.513 | -1.203 | 0.179 | 0.138 |

In north facing triplefins (Figure 3a), the difference between hatting treatments depended on the stimulus presented, as shown by the significant interaction term (Table 2b.1). Shaded individuals stayed significantly closer to a scorpionfish than the averaged controls (Table 2b.2). This effect was absent when exposed to a stone (LMEM stone: hat treatment *p* = 0.097). In south facing triplefins (Figure 3b), shaded individuals did not differ from controls in the distances they kept from either stimulus (Table 2c). Instead, all treatments kept a much larger distance from the scorpionfish than in north-facing triplefins. Because data were collected more than 12 h after adding the fish to the tanks, it was not possible to infer whether south facing triplefins were generally better at detecting a scorpionfish, or whether they had moved further away once they detected its presence. The predictor *time of day* did not contribute significantly to the model, indicating that triplefins had reached a stable distance to the stimulus when observations started.

**Figure 3.**
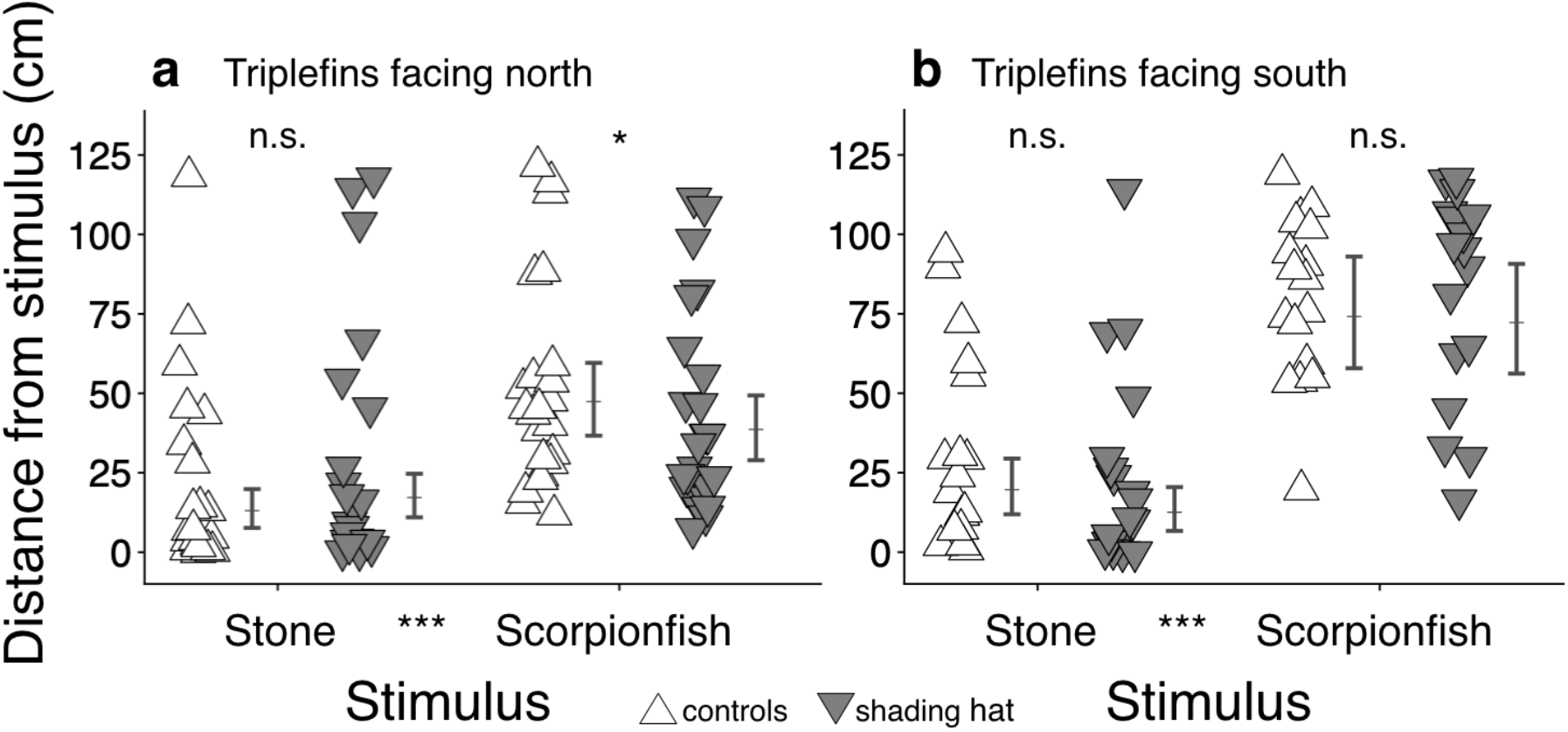
Consequences of hatting in the field at 15 m depth shown as the average distance from the stimulus compartment as a function of stimulus type (stone or scorpionfish), hat treatment, and orientation. **a.** Among north-facing triplefins shaded individuals stayed closer to a scorpionfish than the controls averaged per-triplet (*n* = 24 triplets). **b.** Among south-facing triplefins such effect was absent (*n* = 19 triplets). Symbols: average of 3 measurements per individual; error bars: model-predicted means ± 95 % credible intervals. * = *p* < 0.05, n.s. = *p* > 0.05 (see text and Materials and Methods for details). Note that statistical comparisons between treatments rested on the connected measures *within* triplets and 5 observations per stimulus, making group means and error bars imprecise indicators of the statistical significance of paired measures.

### Rapid short-term changes in distance to a scorpionfish in the field

The previous two experiments observed triplefins only after they had been given much time to inspect the new environment in the tank. To better understand how the resultant distances to the scorpionfish arose, we carried out a follow-up experiment. In contrast to the previous experiments, we assessed the initial response of single, clear-hatted or shaded individuals to a scorpionfish immediately following release in the tank. Triplefin positions were recorded at 7 time points from 1 min until ca. 100 min after release in 10 tanks, again oriented either north or south. Upon careful release at the midpoint of the compartment (25 cm), most triplefins swam towards the display compartment (Figure 4). One min after release, 27 out of 80 fish had approached the scorpionfish to within 7 cm, which is the mean average detection distance estimated by visual modelling for north-facing (6 cm) and south-facing (8 cm) triplefins (see below). Out of these 27, 18 were shading hatted, 9 clear-hatted. This followed from the significant difference in distance to the scorpionfish between treatments illustrated by the non-overlapping 95% credible intervals (Figure 4, Tab. 3). During the following 90-100 min, clear-hatted fish retreated to the opposite half of the tank about 20 min earlier than shading-hatted fish (based on the time at which the curves in Figure 4 cross 25 cm). Both treatments reached a similar equilibrium distance after ~50 min. Tank orientation had no effect, but this may have been a consequence of the shorter distance (50 cm) available to triplefins to move away from the stimulus relative to the previous experiment (125 cm).

**Table 3.**
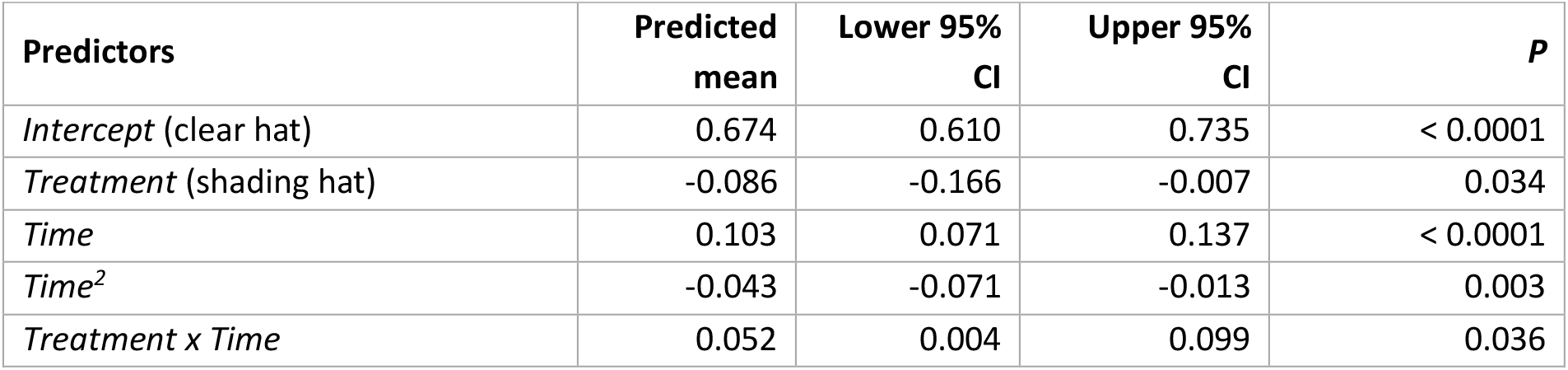
Statistical analysis of the field data presented in Figure 4. Generalized Linear Mixed Model (*n* clear hat = 42, *n* shading hat = 38, *R*^2^_marg_ = 0.46) with proportional distance to the visual stimulus (scorpionfish only) as the response variable. Note that predicted means and their CI are based on a beta distribution with logit link (see Materials and Methods). CI = credible interval. For factorial predictors, estimates are computed using the indicated intercept levels as reference. This choice is arbitrary and does not affect the overall conclusions. This model includes a first-order autoregressive (AR1 = 0.86) variance structure to correct for temporal dependency in the observations of the same individuals.

| Predictors | Predicted mean | Lower 95% CI | Upper 95% CI | *P* |
| --- | --- | --- | --- | --- |
| *Intercept* (clear hat) | 0.674 | 0.610 | 0.735 | < 0.0001 |
| *Treatment* (shading hat) | -0.086 | -0.166 | -0.007 | 0.034 |
| *Time* | 0.103 | 0.071 | 0.137 | < 0.0001 |
| *Time*<sup>2</sup> | -0.043 | -0.071 | -0.013 | 0.003 |
| *Treatment x Time* | 0.052 | 0.004 | 0.099 | 0.036 |

**Figure 4.**
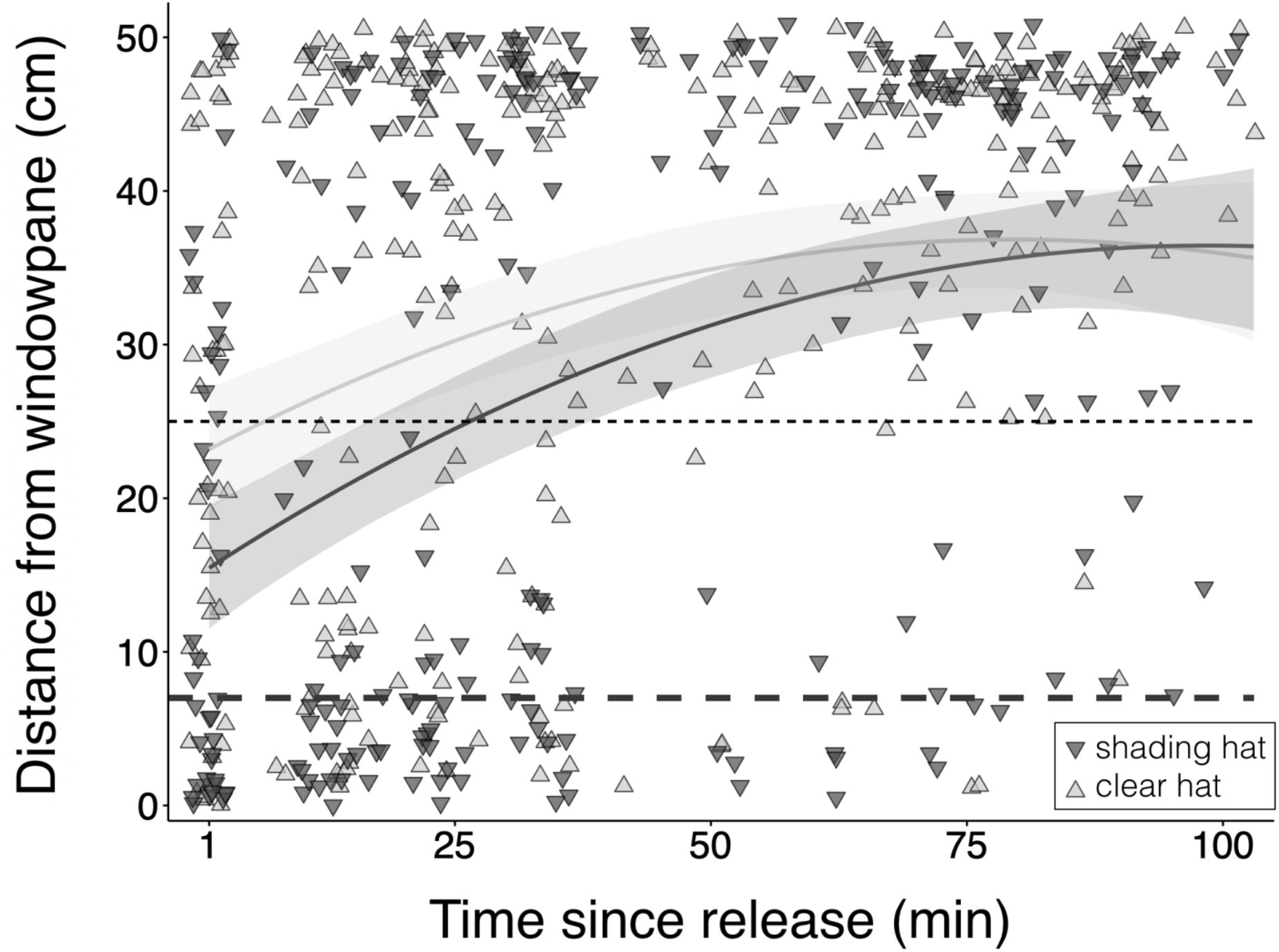
Short-term changes in distance between hatted triplefins and a scorpionfish as a function of time and hat treatment. The first measurement took place about one minute after releasing a single triplefin in the middle of a 50 cm long tank at 10 m depth in the field (*n* clear hat = 42, *n* shading hat = 38). The curved lines show predictions from the most parsimonious, Generalized Linear Mixed Model that describes the movement of shaded (dark gray) and clear-hatted (light grey) triplefins with 95% credible intervals (shaded areas). See Table 3 for statistical details. Each triplefin was observed at 7 time points. Black dashed line: point of release (25 cm). Long-dashed line at 7 cm: average detection distance at which diurnal active photolocation allows a triplefin to induce and perceive scorpionfish eyeshine using a spark, according to visual modelling (Figure 5). Symbols were slightly jittered to reveal overlapping observations in the graph.

### Visual modelling of scorpionfish detectability through induced eyeshine

To validate our experimental results, we implemented visual models to compute the contrast change in the pupil of a scorpionfish perceived by an untreated triplefin when producing an ocular spark. Even when not illuminated by an ocular spark, the pupil of a scorpionfish shows a certain brightness, which improves pupil concealment [9]. This baseline pupil brightness varies with the degree of shading and the substrate on which the scorpionfish sits. Here, we limit ourselves to parameters that match the light conditions of the second field experiment at 10 m and focus on modelling the effect of blue ocular sparks (Figure 1a, b; see [4] for spark types). Relative to a white standard, blue ocular sparks have an average reflectance of 1.34 over the 400-700 nm range, with a maximum average of 2.15 at 472 nm, illustrating the focusing effect of the lens [4]. Further parameters included spectrophotometric measures of the ambient light in the field tanks, scorpionfish pupil size, baseline pupil radiance (Figure 1d), the reflective properties of the pupil and the iris [9], and the triplefin visual system [17–19]. We used the receptor-noise model [20] for estimating chromatic contrasts and Michelson contrasts using cone-catch values of the double cones for achromatic contrasts.

While ocular sparks did not generate chromatic contrast above the discriminability threshold at any distance between the triplefin and the scorpionfish, achromatic Michelson contrasts exceeded the detection thresholds across a broad range of conditions (Figure 5). For comparison, identical calculations for spark-generated contrast changes in a scorpionfish’s iris rather than its pupil showed no perceptible effect under any of the tested conditions. This confirms that subocular light emission is too weak to generate detectable contrasts in structures other than strong directional reflectors such as retroreflective eyes. For north-facing triplefins, the reflection of the ocular spark from a scorpionfish’s pupil would be detectable up to 6 cm under average conditions, increasing up to 10 cm for higher values of ocular spark radiance and scorpionfish eye retroreflectance. Estimated detection distances increased by 2-3 cm for south-facing triplefins.

**Figure 5.**
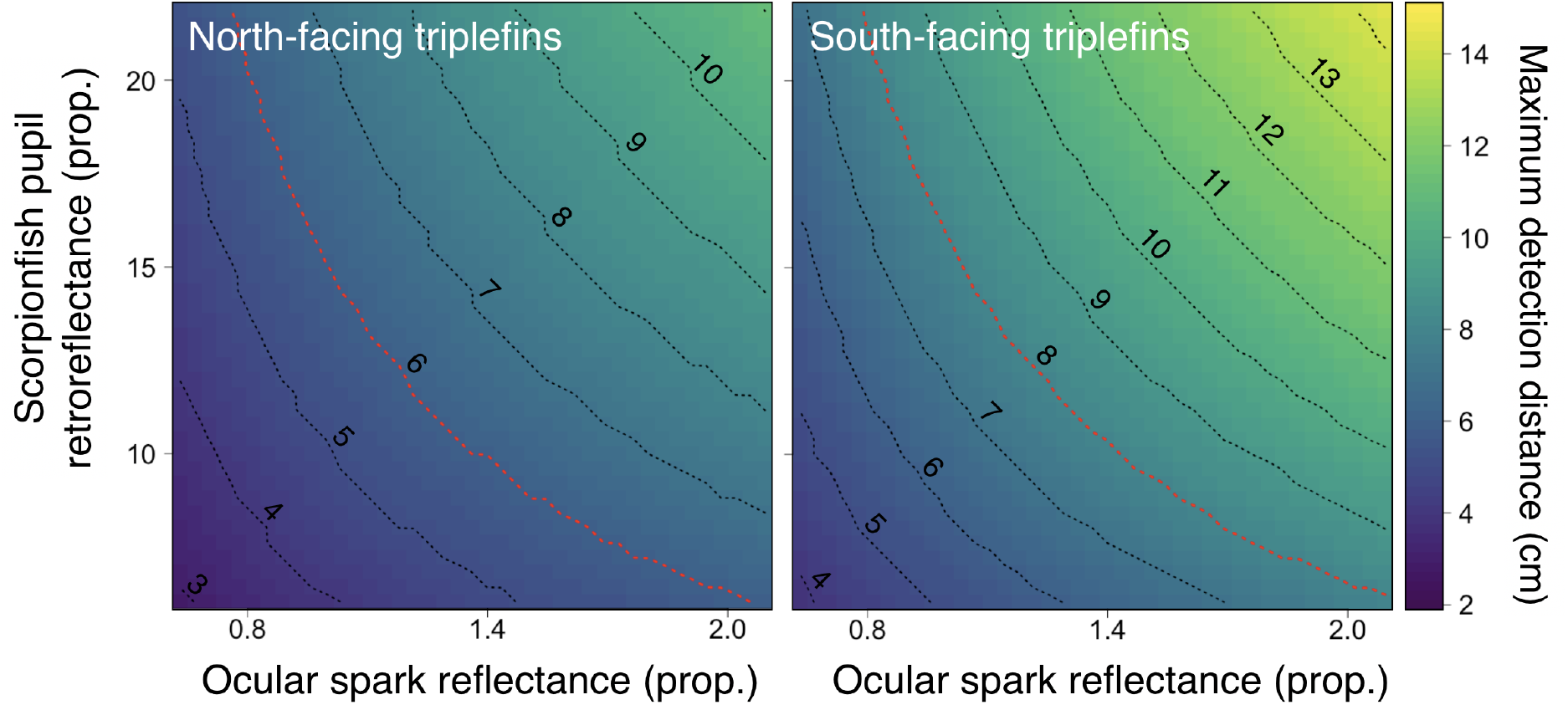
Theoretical detection distances by a triplefin of reflections in a scorpionfish’s eye induced by a triplefin’s blue ocular spark. Visual modelling output using parameters from the field experiment in 10 m, showing maximum detection distance (color, dotted lines) of achromatic contrast differences in a scorpionfish’s pupil as triggered by a triplefin’s blue ocular spark at 10 m depth. The outcome is shown in color as a function of ocular spark reflectance and scorpionfish pupil retroreflectance, separated for north facing and south facing orientations (see Figure 1g-h). The red dotted lines represent intermediate detection distances for both orientations and were summarized as the average detection distance at 7 cm in Figure 4. Values were obtained from calculating the Michelson contrast based on triplefin cone-catches of the double cones for each millimeter distance between 1 and 15 cm, and identifying the maximum distance at which the contrast reached the achromatic contrast threshold of *T. delaisi* (0.8 % [18]). The X and Y axes cover the range of measured values for these predictors (Material and Methods).

## Discussion

We provided a first proof of principle for the diurnal active photolocation hypothesis. Triplefins that were prevented from redirecting light with their iris kept shorter distances from a scorpionfish than control-treated individuals. Visual modelling confirmed that this can be explained by diurnal active photolocation: triplefins can induce a perceptible contrast in a scorpionfish’ retroreflective pupil over biologically relevant distances. We conclude that controlled light redirection can improve visual detection substantially under realistic conditions. At the same time, it is important to stress that diurnal active photolocation is not failproof. Visual modelling also set detection limits of this mechanism. For unfavorable, yet realistic parameter values, the explored parameter space predicts detection distances so short that they are likely to fall within the striking range of a cryptic sit-and-wait predator such as a scorpionfish [21–23].

### Observed distances versus detection distances

Visual modelling predicted shorter distances over which diurnal active photolocation can improve predator detection than the actual triplefin-scorpionfish distances observed in the laboratory and first field experiment (Figures 2 and 3). This discrepancy arose because the first two datasets did not measure the distance of detection or closest approach, but the distance established after acclimatization to the new environment. The second field experiment (Figure 4) complemented these observations by showing that immediately after release in a new environment, many triplefins moved towards the display compartment, resulting in closer distances to the scorpionfish in shaded than in clear-hatted triplefins. Although such data are not available for the first two experiments, we assume that a similar initial assessment of the display compartment explained the differences between treatments still visible the next day.

### Alternative approaches to test one question

The fact that the first field experiment showed weaker treatment effects than the similar laboratory experiment illustrates the importance of replicating this type of experiment under laboratory as well as natural light fields, even if this involves new challenges such as temporal variation in sunlight, and distraction by fish naturally occurring around the tanks. Hence, the power of this study lies in the demonstration of a similar treatment effect confirmed across three independent and different experiments. The statistical power of the first two experiments was enhanced by treatment comparison within triplets (*triplet* as random factor) to compensate for triplet-specific variation. Shaded fish may, however, have followed control-treated fish, weakening a treatment effect. In the second field experiment, which was designed to test for immediate responses in single individuals, we found a treatment effect that was qualitatively similar to the one observed in the first two experiments. This suggests that the results were not strongly affected by inter-individual interactions.

### Possible artifacts caused by hatting

The superglue used to attach the hats is commonly used in veterinarian surgery, including fish. The data presented here, as well as additional experiments [24] showed that hatting did not appear to affect triplefin behavior, except for scorpionfish detection. Yet, it is still conceivable that a shading hat reduced a triplefin’s visual field, offering an alternative explanation to poorer detection of a scorpionfish. Hat design, however, anticipated this problem. As explained in Materials and Methods, hats were folded as small “umbrellas”, hovering well above the fish’s eyes (Figure 1b-c). Consequently, the forward viewing angle was well above 45° from horizontal. Moreover, triplefins typically sit in an upright position propped up on their pectoral fins. Given that the visual cues were presented at the same level as the triplefins, we have therefore no doubt that both stimuli fell well within the viewing range (scorpionfish eye < 4 cm above the substrate). Moreover, if hats would have blocked the forward view, even clear-hatted fish must have seen the world in a distorted way when looking through their hat from a slanted angle. Yet, there was no difference between the clear-hatted and unhatted control. The response of all treatments to the safe stimulus *stone* did not differ, once more indicating that overall forward vision did not appear to be affected. We therefore consider it safe to exclude visual obstruction by a hat as an alternative explanation for the experimental results.

### Ubiquity of ocular light redirection and its consequences

We assume that the iridophore patch on the lower iris of a triplefin is a diffuse, Lambertian reflector, which has been confirmed for the equatorial plane [25]. It may therefore function as a short-distance detector that covers most of the hemispherical zone seen by a single eye, and that is effective over short distances only. In lantern and flashlight fish, subocular light organs are also diffuse sources [1, 2]. Many other fish, however, possess silvery irides with near-specular properties. Such reflectors are more directional, possibly allowing specific illumination of the scene or objects of interest over larger distances. Specular reflection, however, may increase visibility to others, including predators. This trade-off may explain the variation seen in types of ocular light redirection in diurnal fish families, which varies from highly conspicuous to very subtle as in triplefins [4, 26]. In the target organisms, highly reflective structures such as retroreflective eyes [9, 14] or reflectors in cryptic crustacean prey [4, 25] are also common and diverse. For now, it is too early to speculate which structures and conditions may allow active photolocation in other species because quantitative measurements and experimental data are still missing. Yet, it is clear that the basic building blocks required for diurnal active photolocation are ubiquitous. One may even postulate that the properties of well-camouflaged, cryptic predators are partly explained as an evolutionary response to the use of diurnal active photolocation by their prey. Most marine cryptobenthic predators indeed show eye adaptations that hamper their discovery. Stonefish (*Synanceia*) and frogfish (*Antennarius*) have surprisingly small eyes for their body size. Other species have skin flaps that partially cover the pupil as in crocodile fish (*Papilloculiceps*) and some scorpionfishes (some *Scorpaenopsis species*), or possess slit-like pupils as in some flatheads (*Thysanophrys*), flounders (*Bothus*) and sandperches (*Parapercis*). In lionfish (*Pterois*) the eyes are included in one of several black vertical lines on the body. All these traits reduce pupil size, distort its shape or mask its presence. Since eyes are commonly used for face recognition [27, 28] such modifications make detection harder for prey to detect a cryptic predator [29] or for a predator to detect cryptic prey [30]. Although all this can be explained by unaided vision alone, it is a tantalizing possibility that diurnal active photolocation is also involved. A special feature of scorpionfish in this context is their diurnal eyeshine, resulting in an unusually “bright”, not black pupil caused by a combination of light reflection and transmission [14, 31]. It improves camouflage by reducing the contrast between a pupil and the surrounding skin. It represents an alternative mechanism to impair visual detection, and also reduces the effectiveness of active photolocation. At the same time, however, the retroreflective component of diurnal eyeshine in scorpionfish can be exploited by fish with subocular light redirection, as shown here for triplefins. Daytime eyeshine is present in other cryptobenthic predators, particularly in species such as devilfishes (*Inimicus*), toadfishes (*Halophyme*) and seem ubiquitous in scorpionfishes (*Scorpaena, Scorpaenopsis, Rhinopias, Pteroidichthys*). In almost all of these, pupils are large and circular, suggesting that daytime eyeshine relaxes the need to mask the shape or size of a pupil. The selective forces on cryptobenthic predators generated by active photolocation are identical to those that can be expected from regular visual detection alone, which is why experimental manipulation is required to separate their role.

### Future perspectives

This study represents an important first step towards our understanding of a complex visual interaction between a cryptic predator and its visual prey. Topics for future work involve a manipulation of properties such as the baseline radiance and retroreflectance of scorpionfish eyes, or ocular spark size and brightness in triplefins. Furthermore, we see a potential for tests in fish species with silvery, more specular irides or other forms of light redirection. Targets other than predators are also promising. There is some indirect evidence for prey detection using active photolocation [25, 26, 32], but more empirical studies are needed for confirmation. In addition, we described ocular sparks as illuminants, but this does not preclude other functions such as intraspecific communication [4] as proposed for subocular light organs in flashlight fish [2]. Triplefins, however, have a rich signaling repertoire involving body postures and fin raising or flicking. How ocular sparks fit in is unclear and remains to be studied. A role in inter-specific signaling is also conceivable: assuming ocular sparks represent a signal to attract the attention of a scorpionfish, it may respond by turning its gaze towards the triplefin. If so, it would improve the efficiency of active photolocation because retroreflection of a lens eye is strongest when it is focused on the target’s light source. Finally, this work made us realize that surprisingly few credible facts have been published concerning the visual and behavioral interactions between cryptobenthic predatory fish and their fish prey. This field has thus far been governed by intuitive but untested interpretations and therefore offers plentiful opportunities for those prepared to explore it.

## Materials and Methods

### Model species and location

Triplefins (Fam. Tripterygiidae) are small, cryptobenthic micropredators that favor marine hard substrates. Our model species is *Tripterygion delaisi*. With a standard length of 3–5 cm it is one of the larger members of this family. *T. delaisi* occurs in the NE-Atlantic and Mediterranean on rocky substrates between 3-50 m depth, but reaches highest densities in 5-15 m. Aside from breeding males, it is highly cryptic and regularly produces blue and red ocular sparks [4].

*Scorpaena porcus* (Fam. Scorpaenidae) is a cryptobenthic sit-and-wait predator (12–20 cm) from coastal marine hard substrates and seagrass habitats across the NE-Atlantic and Mediterranean Sea [33]. It responds to moving prey; non-moving or dead prey is ignored. Small benthic fish, such as triplefins, are often a component of its diet [34]. It possesses a reflective *stratum argenteum* and partially translucent retinal pigment epithelium that allows the generation of daytime eyeshine, which is considered to improve pupil camouflage [9].

All experiments were conducted in Calvi (Corsica, France) under the general permit of STARESO (Station de Recherches Sous Marines et Océanographiques). The hatting technique was developed at the University of Tübingen under permit ZO1-16 from the Regierungspräsidium Tübingen prior to the field experiments.

### Hatting technique to block ocular sparks

We blocked ocular spark formation by means of mini-hats excised from polyester filter sheets using a laser cutter (RLS 100, AM Laserpoint Deutschland GmbH, Hamburg, Germany). A dark red filter with average transmission 1 % was used as the shading treatment (LEE #787 “Marius Red”, LEE Filters, UK). Clear filter hats (LEE #130, “Clear”) were used in the first control group, and no hat, but the same handling procedure, in the second control group. Hats were individually adjusted with clippers and folded into their final configuration with a triangular base for attachment and raised, forward-projecting wings to shade the eyes from downwelling light only. Hats formed an “umbrella” well above the eye, allowing full eye movement in all directions (Figure 1b-c). They varied from 6 to 9 mm in diameter, matching individual head size. Given that *T. delaisi* possesses a fovea that is looking forward and downward when the eye is in a typical position [17], it seems unlikely that shading alone may have resulted in poorer visual detection of a benthic predator in front of the fish relative to a triplefin without hat and without ocular spark. Animals in the clear-hatted and unhatted control groups regularly generated ocular sparks both in the laboratory and in the field.

Triplefins were collected using hand nets while SCUBA diving and brought to a stock aquarium in the laboratory. Individuals were anaesthetized (100 mg L^−1^ MS-222 in seawater, pH = 8.2) until all movements ceased except for breathing (3–4.5 min). Subsequently, the dorsal head surface was gently dried with paper tissue. Hats were glued to the triangular dorso-posterior head area just behind the eyes using surgical glue (Surgibond, Sutures Limited, UK or Vetbond Tissue Adhesive, 3M). After allowing the glue to polymerize for 45 s, fish were moved into recovery containers with aerated seawater. Individuals regained consciousness and mobility within 5–10 min. This non-invasive hat fixation protocol minimized impacts on the fish’s natural behavior and health, as indicated by a 97.4 % survival rate. As a trade-off, however, hats detached within 0–4 days, which reduced the number of fish that could be used for analysis (see Statistical analysis). All fish were treated and included in trials once, but kept in the laboratory for recovery. They were returned to the field after completion of the experiment.

Pilot experiments confirmed that typical behaviors such as fin flicks, push-ups, active movement across the substrate, and head and eye movements did not differ between shading and control treatments [24].

### Laboratory experiment

Four aquaria (L × W × D: 130 × 50 × 50 cm^3^) were used for 20 experimental runs, each employing a new triplet of size-matched *T. delaisi*. In each tank, we placed a rock and a scorpionfish in two separate perforated containers (L × W × H: 24 × 14 × 16 cm^3^) with a glass front. The bottom of the aquarium was barren (avoided by the fish), except for a 10 cm strip of gravel placed along the long side of the tank, providing a sub-optimal substrate. Each tank was illuminated with a 150 W cold white LED floodlight (TIROLED Hallenleuchte, 150 W, 16000 Lumen) shielded with a LEE Filters #172 Lagoon Blue filter to simulate light at depth. The area of the tank where stimuli were displayed was shaded. Both stimuli were simultaneously present in the tank, but only one was visible on a given day. On day one, all fish were treated and placed in the tank in the evening. Observations took place on days two and three. Two aquaria started with stimulus “scorpionfish”, the other two with “stone”, and stimuli were swapped after day two. Hence, all triplets were exposed to a stimulus for one full day. Since fish are moving regularly, we assessed the distance to the stimulus five times per day, 5 min per individual, at 0800, 1100, 1300, 1500 and 1800.

### Replicate experiment in the field

We replicated the laboratory experiment in the field using ten tanks of spectrally neutral Evotron Plexiglas (L × W × D: 150 × 25 × 50 cm^3^) placed at 15 m depth on a sandy patch in the seagrass meadow in front of STARESO. We used local silica sand mixed with gravel as substrate for the compartment in which triplefins were kept (125 x 25 cm^2^). It was separated from a display compartment (15 x 25 cm^2^) for the shaded visual stimulus with transparent Plexiglass. Another similar-sized compartment behind the display compartment was used to keep the stimulus not currently visible to triplefins, separated by an opaque grey PVC plate. All separators were perforated to assure that a scorpionfish invisible to the triplefins could be chemically perceived even when the stone was visible. Visual contact between tanks was excluded by surrounding each enclosure with 10 cm white side covers along the bottom edge. As a response variable, we noted the distance of each individual from the stimulus compartment three times a day at 0900, 1200 and 1500 for two days following deployment in the early evening of the first day. Stimuli were always changed after the first observation day. Triplets were replaced every three days. In total, 50 triplets were tested.

### Second field experiment: short-term response over time

We carried out a second field experiment with the goal of observing the temporal pattern of triplefin inspection behavior immediately after release. To this end, we only tested shading hatted and clear-hatted triplefins individually (not in pairs or triplets) and exposed them to a shaded scorpionfish only (no stone to maximize sample size). As before, we used 10 Plexiglass tanks, 5 with triplefins facing north, another 5 with triplefins facing south. Tanks were identically built (Figure 1) and equally high, but with a smaller footprint, offering 50 x 25 cm^2^ substrate for the triplefins and 12 x 25 cm^2^ for the scorpionfish. To improve SCUBA diving safety, tanks were positioned at a depth of 10 m and mounted on floats with 4 plastic chains attached to 1 m metal rods anchored in the ground. The substrate on which triplefins were placed was covered with darker sand than in the previous experiments, and we used black side covers to block their view to the outside, creating a slightly darker background than in the previous experiment. Scorpionfish (*n* = 10) were kept as a resident in the display compartment. One triplefins was added to each tank at the beginning of a dive and its position determined about 1 min after release. Once all triplefins had been released and their distance recorded for the first time, each tank was visited another 3 times during this first dive. After a ~30 min surface interval, the divers went back to collect another 3 data points, after which all triplefins were removed. Due to this procedure, time intervals between tanks and surface interval between first and second dive varied slightly. Eight cohorts of 10 triplefins were observed, 38 shaded and 42 clear-hatted triplefins. Using controlled randomization, treatments were equally distributed across cohorts, tank ID and tank orientations to prevent any systematic bias.

### Statistical analysis

#### Repeatability analysis

In all three experiments, distance measurements were not blind for hat treatment. However, room for error was limited as we did not interpret a behavior, but merely noted the position of the head of a fish relative to a ruler placed alongside the tank. In the laboratory, fish and ruler were very close to each other and therefore easy to align to take virtually error-free measurements. In the field, the SCUBA diver was hovering above the tank and used rulers on both long sides for alignment and to determine fish position. To test repeatability in the field, the two divers who collected the distance data in the field (MS, UKH) determined 116 distances of triplefins in the 15 m field tanks. Using the R package *rptR* [35], datatype *Gaussian* and 1000 permutations, the repeatability estimate was *R* = 0.995 (Likelihood Ratio Test: *P* < 0.0001).

#### Statistical model choice and pooling of controls

Behavioral data were analyzed using Generalized Linear Mixed Models (GLMM) with the lme4 package [36] and glmmTMB package [37] for R v3.4.3. [38]. For the first two experiments, we first compared the two control treatments (sham and clear hat) to verify that hatting a fish did not affect behavior, and to confirm their ability to distinguish a cryptic predator from a stone. Because controls did not differ, we then averaged the data of the two control-treated fish per triplet per observation for the final models and compared them to the shaded treatment. This allowed us to also include triplets in which only the clear-hatted fish had lost its hat for the comparison with the shaded fish (such triplets had been excluded from the comparison of the controls). This explains the variation in triplet numbers in the final analyses. Distance from the display compartment was used as the response variable in all three models, implemented using a normal distribution for the first two experiments and a beta binomial distribution (link = log) for the third one.

#### Predictors and transformations

For the laboratory experiment, the initial fixed model component included the main predictors *stimulus* (stone vs scorpionfish), *hat treatment* (no hat vs clear hat, or averaged controls vs shaded) and their interaction. We further included the fixed covariates *time of day* for each observation, *stimulus order, cohort and tank ID*. The models for the replicated field experiment were identical, but also included the fixed factor *orientation* (north or south) and its interactions with the main predictors. We square-root-transformed the response variable *distance* to improve residual homogeneity in the analysis of the first field experiment. The transformation of the response variable did not cause any change in the effects of the interactions between covariates. Models to compare the response of controls vs shaded fish were calculated separately for north vs south orientation because fish responded differently to the scorpionfish depending on orientation (Figure 3, Table 2).

For the third experiment, the initial fixed model component included the main predictors *hat treatment* (clear hat or shaded), time, orientation and their three-way interaction. We also included time as a quadratic component to explain the non-linear patterns of the data, assessed using the *gam* function of the mgvc R package [39], and the covariate day, as data were collected on three subsequent days. The response variable was transformed as proportion (0 < x < 1) of distance obtained by dividing all distances by the maximum length of the tank plus one (51 cm). The transformation of the response variable did not affect the interactions between covariates, yet allowed us to implement a beta binomial distribution, thus improving residual homogeneity. We finally included a first-order autoregressive (AR1) variance structure to correct for temporal dependency in the observations of the same individuals.

#### Triplet as random factor and model selection

In the first two models, the initial random component contained triplet ID with random slopes over the hat treatment. This accounts for the repeated measurements of each triplet and captures variation arising from different hat-treatment responses among triplets [40]. Random slopes were uninformative and subsequently removed. In the third model, the random component included triplefin ID, tank ID and cohort. We then performed backward model selection using the Akaike Information Criterion (AIC) to identify the best-fitting model with the smallest number of covariates [41]. We only report the reduced final models and provide proxies for their overall goodness-of-fit (marginal and conditional *R*^2^) using piecewiseSEM [42]. The marginal *R*^2^ expresses the proportion of variation explained by the model considering fixed factors only, whereas the conditional *R*^2^ expresses the same including the random factors [43]. We used Wald z-tests to assess the significance of fixed effects. To explore significant interactions between stimulus and hat treatment, we implemented new models within the two levels of the stimulus treatment. Model assumptions were validated by plotting residuals versus fitted values and each covariate present in the full, nonreduced model [44].

### Estimating scorpionfish pupil radiance with and without ocular spark

We assumed both triplefins and scorpionfish were looking orthogonally at one another to calculate the photon flux of the scorpionfish pupil reaching the triplefin pupil (SI 1). Using retinal quantum catch estimates, we calculated the chromatic contrast [20] between the scorpionfish pupil with and without the contribution of the blue ocular sparks. The achromatic contrast between the same two conditions was estimated by calculating the Michelson contrast using the quantum catches of the two-long-wavelength photoreceptors. For comparison, we also performed the same calculations using photon flux from the scorpionfish iris with and without the contribution of an ocular spark. We parameterized the equations using measurements of: (1) ambient light in the tanks at 10 m depth, (2) the range of ocular spark radiance under downwelling light conditions, (3) baseline scorpionfish pupil radiance in the experimental tanks, (4) sizes of triplefin pupil, ocular spark and scorpionfish pupil, and (5) scorpionfish pupil and iris reflectance [9].

Spectroradiometric measurements were obtained with a calibrated SpectraScan PR-740 (Photo Research, New York USA) encased in an underwater housing (BS Kinetics, Germany). This device measures spectral radiance (watts sr^−1^ m^−2^ nm^−1^) of an area with defined solid angle. The downwelling light was estimated by measuring the radiance of a polytetrafluoroethylene (PTFE) diffuse white reflectance standard (Berghof Fluoroplastic Technology GmbH, Germany) positioned parallel to the water surface from a 45° angle. Radiance values were subsequently transformed into photon radiance (photons s^−1^ sr^−1^ m^−2^ nm^−1^).

We determined the relationship between the radiance of the ocular spark and that of a white PTFE standard exposed to downwelling light in live triplefins. Fish mildly sedated with clove oil (*n* = 10) were placed in an aquarium illuminated with a Leica EL 6000 source and a liquid light guide suspended ~20 cm above the tank. Spark radiance was normalized by comparing it to a white standard at 45° from normal positioned at the same location as the fish. For each fish, three measurements were obtained from each eye. The highest value for each fish relative to the standard was used for the model. The sizes of the triplefin pupil (*n* = 35), the ocular spark (*n* = 10), and the scorpionfish pupil (*n* = 20) were measured in ImageJ [45] using scaled images. Natural baseline pupil radiance of three different scorpionfish was measured orthogonally to the pupil from the perspective of the triplefins during the field experimental trials using a Photo Research PR-740 spectroradiometer.

Solid angles of the ocular spark as perceived from the perspective of the scorpionfish, and the pupil of the scorpionfish as perceived by the triplefin were computed using simple calculations (see below).

### Visual models and maximum detection distance

The receptor-noise limited model for calculation of chromatic contrast was informed using triplefin ocular media transmission values, photoreceptor sensitivity curves [19, 46], and the relative photoreceptor density of single to double cone of 1:4:4 as found in the triplefin fovea [17]. We used a Weber fraction (*ω*) value of 0.05 as in previous studies [47, 48]. Chromatic contrasts are measured as just-noticeable differences (JNDs), where values greater than 1 are considered to be larger than the minimum discernible difference between two objects. We calculated the Michelson achromatic contrast as

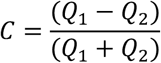

where *Q*_1_ and *Q*_2_ are the quantum catches of the two members of the double cones which are associated with the achromatic channel, under photon flux_1_ and photon flux_2_. Flux_1_ is the sum of the photon flux into a triplefin’s eye caused by the baseline radiance of a scorpionfish pupil and the photon flux caused by the retroreflection of an ocular spark in the scorpionfish pupil (sum of equations (2) and (6) below). Flux_2_ is calculated from the baseline radiance of a scorpionfish pupil only (no ocular spark reflection, equation (2) below). We determined the maximum discernible distance of the ocular spark radiance reflected through a scorpionfish pupil by calculating the chromatic and achromatic contrast at each millimeter, between 1 and 15 cm, and extracting the first value at which the contrast was equal to or exceeded the threshold of 1.0 JND for chromatic contrasts and 0.008 for Michelson contrasts as measured in *T. delaisi* [18] and other fish species [49]. All visual models were performed using the R package pavo [50].

**Table 4.** Symbols and indices used in the equations to calculate the photon flux of the scorpionfish pupil reaching the triplefin, with and without the contribution of an ocular spark.

| ***Symbol*** | ***Definitions and units*** |
| --- | --- |
| $L$ | Photon radiance (photons $s^{-1} sr^{-1} m^{-2}$) |
| $S$ | Blue ocular spark reflectance (proportion in relation to PTFE white standard) |
| $d$ | Distance between triplefin and scorpionfish (m) |
| $\Delta$ | Mean displacement of ocular spark relative to triplefin pupil center (0.00109 m) |
| $r$ | Radius (m) |
| $\Omega$ | Solid angle (sr) |
| $R$ | Reflectance of coaxially illuminated scorpionfish pupil (prop. in relation to PTFE white standard) |
| $\kappa$ | Diffuse attenuation coefficient ($m^{-1}$) |
| $\Phi$ | Photon flux (photons $s^{-1}$) |
| ***Indices*** | ***Meaning and use*** |
| $0$ | Distance = 0, as used in $L_0$ |
| $w$ | Used for downwelling light from the water surface, used in $L_w$ |
| $ns$ | Abbreviation for "no ocular spark", used in $\Phi_{ns}$ |
| $os$ | Abbreviation for ocular spark of a triplefin, used in $L_{os}$, $r_{os}$ and $\Omega_{os}$ |
| $sp$ | Abbreviation for scorpionfish pupil used in $L_{sp}$, $r_{sp}$ and $\Omega_{sp}$ |
| $t$ | Abbreviation for triplefin, used in $r_t$ |

### Visual model details

#### Triplefin – scorpionfish interaction

The starting conditions assume that both fish look at each other at normal incidence, i.e. the full area of the pupil of the triplefin is visible to the scorpionfish and vice versa. Solid angles are computed as explained below, assuming the ocular spark is positioned at the edge of the iris (displacement from pupil center *Δ* = 1.09 mm) in the plane of the triplefin pupil.

#### Photon flux without ocular spark

The photon radiance of the scorpionfish pupil reaching the triplefin (*L_d_*) is a function of the measured scorpionfish pupil photon radiance (*L*_0_) attenuated by the aquatic medium over distance *d* such that

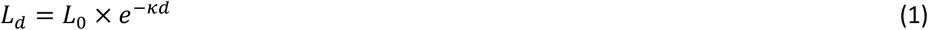

The photon flux reaching the retina of the triplefin without the ocular spark (*Φ_ns_*) (Figure S1) is the proportion of attenuated photon radiance reaching the triplefin’s pupil (*L_d_*) multiplied by the solid angle of the scorpionfish pupil (*Ω_sp_*) and the area of the triplefin pupil 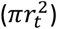:

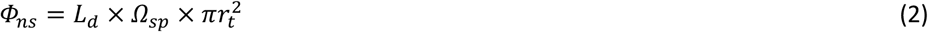

This value was used to calculate the quantum catches *Q*_1_ and *Q*_2_ mentioned earlier.

#### Photon flux with ocular spark

The photon radiance of the ocular spark reaching the scorpionfish (*L_os_*) is a function of the radiance of a PTFE white standard parallel to the water surface (*L_w_*), the focusing power of the lens, and the reflective properties of the iridal chromatophores on which the light is focused. For now, the focusing power and reflective properties have only been measured together as blue ocular spark reflectance (*S*) relative to *L_w_*:

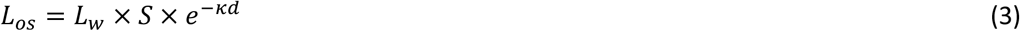

The radiance of the scorpionfish pupil (*L_sp_*) defined as the proportion of the attenuated ocular spark photon radiance that reaches the scorpionfish pupil and is re-emitted towards the triplefin is estimated by multiplying the photon radiance of the ocular spark reaching the scorpionfish (*L_os_*) with the solid angle of the ocular spark as seen by the scorpionfish (*Ω_os_*) and the retroreflectance of the scorpionfish pupil with illumination co-axial to the receiver (*R*). Because the properties of the retroreflective eye are measured in relation to a diffuse white standard, the photon exitance from the scorpionfish pupil is converted to photon radiance by dividing by *π* steradians:

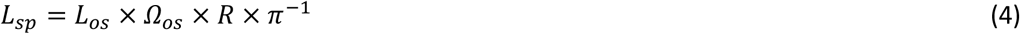

The scorpionfish pupil radiance (*L_sp_*) travelling towards the triplefin pupil is further attenuated, and the photon flux reaching the triplefin’s retina (*Φ_os_*) is obtained by multiplying the attenuated radiance by the solid angle of the scorpionfish pupil, and the area of the triplefin pupil:

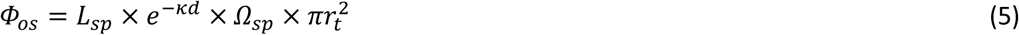

The photon flux generated by the ocular spark, which reaches the triplefin retina after being reflected by the scorpionfish pupil is therefore approximated by (see also Figure S2):

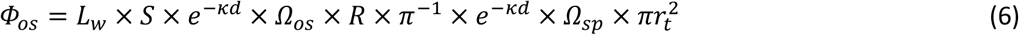

The total photon flux reaching the retina of the triplefin with the ocular spark is then the sum of equations (2) and (6) (Figs. 5 and 6 combined). This sum was used to calculate the quantum catches *Q*_1_ and *Q*_2_ from a scorpionfish eye illuminated by an ocular spark, as mentioned earlier.

**Figure 6.**
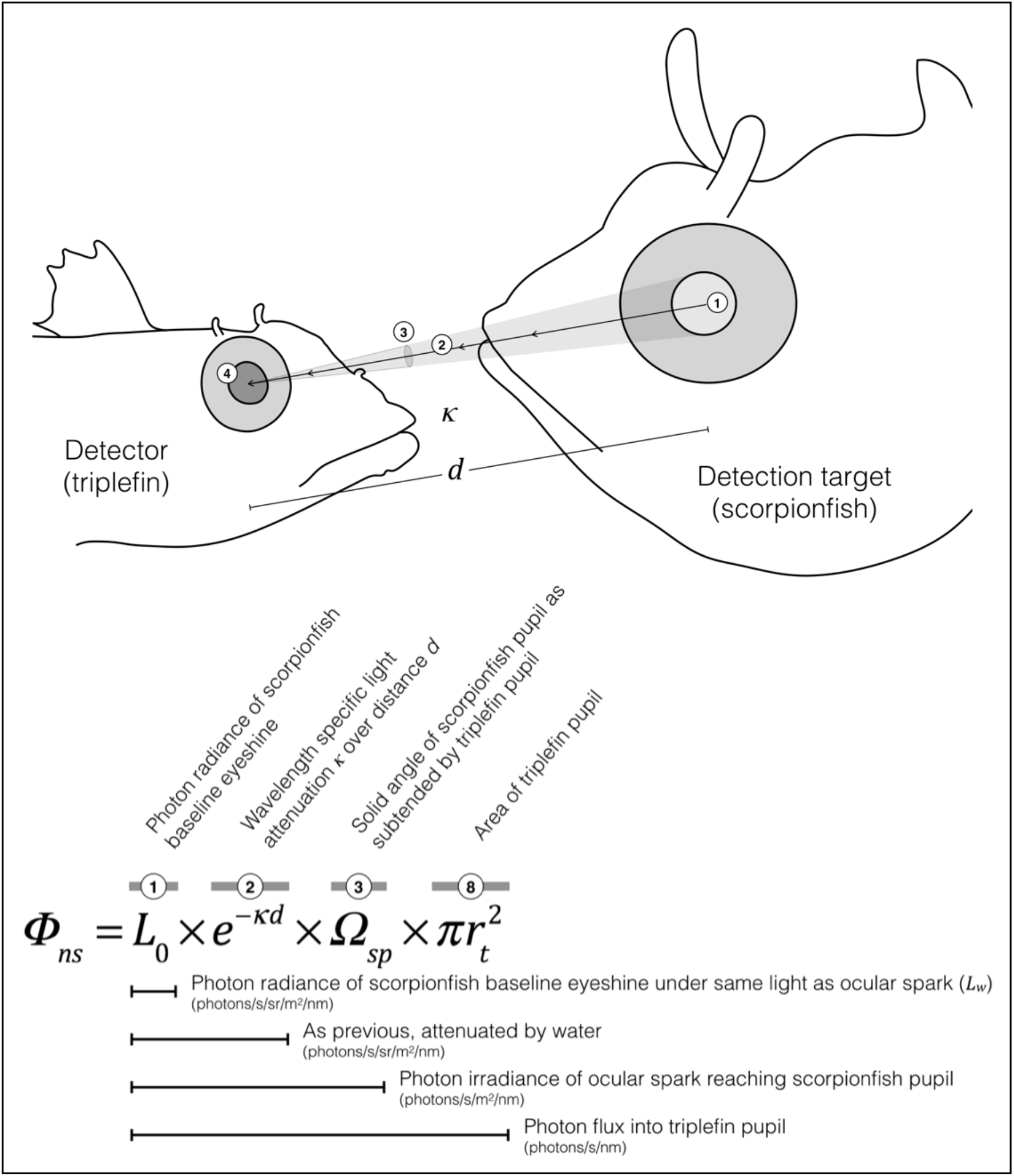
Visual representation of how the photon flux ***Φ***_*ns*_ originating from baseline scorpionfish eyeshine entering a triplefin’s pupil is calculated. This case excludes the effect of an ocular spark, which is shown in Figure 7.

**Figure 7.**
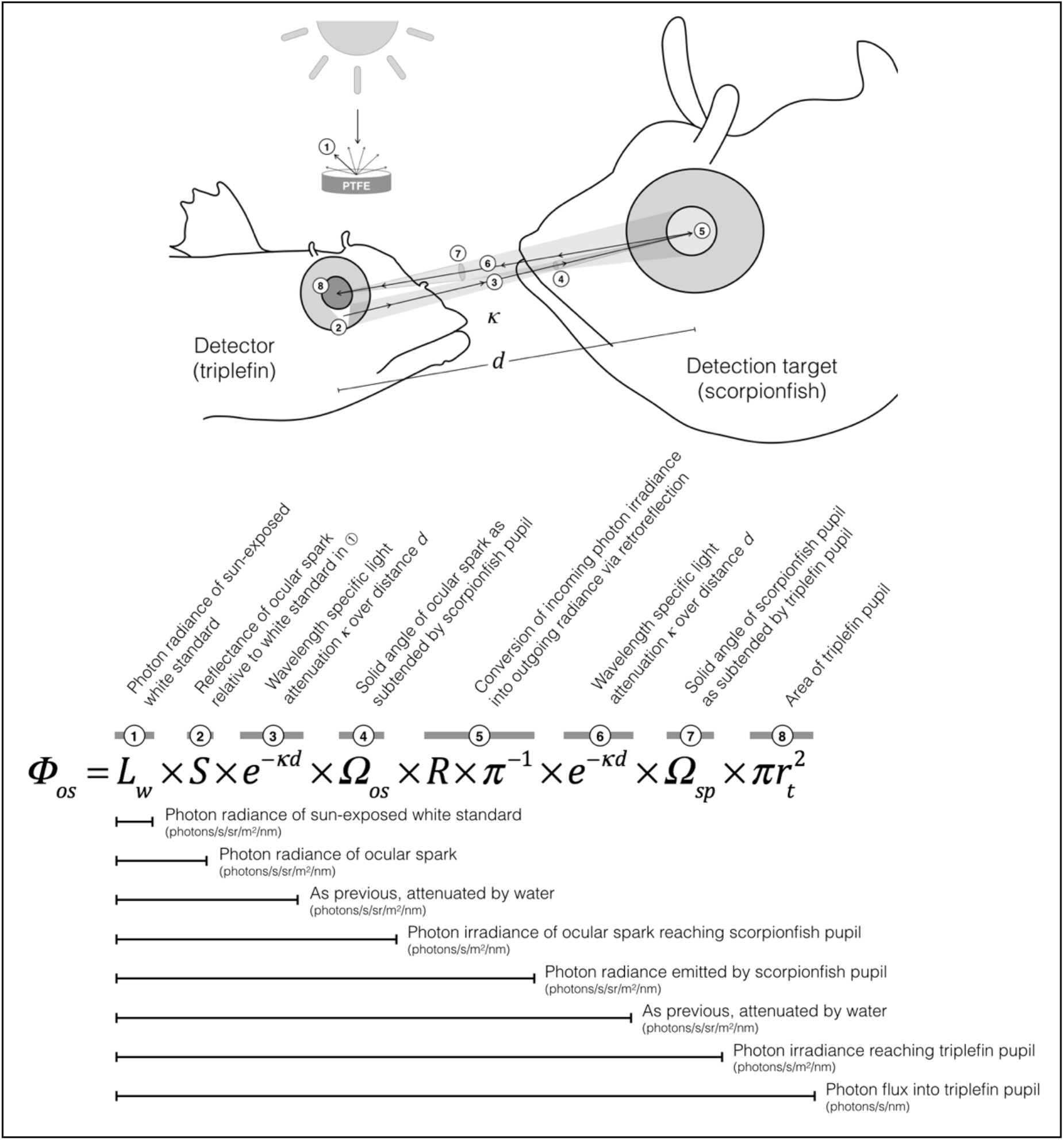
Visual representation of how much of the photon flux ***Φ***_*os*_ generated by a triplefin’s ocular spark is reflected as scorpionfish eyeshine and ultimately reaches a triplefin’s pupil. This effect needs to be added on top of baseline scorpionfish eyeshine (explained in Figure S1), to obtain the total photon flux from a scorpionfish eye reaching the eye of a triplefin with its ocular spark on.

#### Calculation of solid angles

The solid angle of the scorpionfish pupil (*Ω_sp_*) as perceived by the (dimensionless) center of the triplefin’s pupil at distance *d* was estimated using the formula

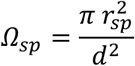

The solid angle of the ocular spark as seen from the perspective of a scorpionfish eye (*Ω_os_*) needs to be corrected for the fact that the ocular spark is below the triplefin’s pupil by a distance *Δ* = 0.00109 m. The radius of the ocular spark at this distance as perceived by the scorpionfish can be calculated by multiplying the original diameter *r*_os_ with the ratio of the original distance *d* divided by the hypotenuse of the right-angled triangle defined by *Δ* and *d*:

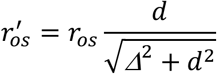

The solid angle of the ocular park as perceived by the (dimensionless) center of the scorpionfish’s pupil can then be calculated as

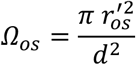

## Acknowledgments

Thanks to Martin J. How for useful suggestions on an earlier draft. Jonas Dornbach, Thomas Griessler, Katharina Hiemer, Michael Karcz, Valentina Richter, Peter Tung, Sabine Urban, Laura Warmuth and Florian Wehrberger supported data collection in the field. Gregor Schulte provided creative and technical support. Thanks to Pierre Lejeune, director of STARESO and his staff for providing excellent working conditions. N.K.M. was supported by Koselleck Grant Mi 482/13-1 from the Deutsche Forschungsgemeinschaft and Experiment! grant Az. 89148 and Az. 91816 from the Volkswagen Foundation. P-P.B. was funded by a Postdoctoral Fellowship from the Natural Sciences and Engineering Research Council of Canada.

